# A mysterious cloak: the peptidoglycan layer of algal and plant plastids

**DOI:** 10.1101/2023.05.02.539068

**Authors:** Alexander I. MacLeod, Michael R. Knopp, Sven B. Gould

## Abstract

The plastids of algae and plants originated on a single occasion from an endosymbiotic cyanobacterium at least a billion years ago. Despite the divergent evolution that characterizes the plastids of different lineages, many traits such as membrane organisation and means of fission are universal – they pay tribute to the cyanobacterial origin of the organelle. For one such trait, the peptidoglycan (PG) layer, the situation is more complicated, and little is known about its distribution and molecular relevance in green algae and land plants. Here, we investigate the extent of PG presence across the Chloroplastida using a phylogenomic approach. Our data support the view of a PG layer being present in the last common ancestor of land plants and its remarkable conservation across bryophytes that are otherwise characterized by gene loss. In embryophytes, the occurrence of the PG layer biosynthetic toolkit becomes patchier, but the availability of novel genome data questions previous predictions regarding a functional coevolution of the PG layer and the plastid division machinery-associated gene FtsZ3. Furthermore, our data confirm the presence of penicillin-binding proteins (PBPs) in seed plants, which were previously thought to be absent from this clade. The thicker and seemingly unchanged PG layer armouring the plastids of glaucophyte algae might still provide the original function of structural support, but the same can likely not be said about the only recently identified and ultrathin PG layer of bryophyte and tracheophyte plastids. In combination with the apparent lack of some genes thought critical for PG layer biosynthesis in land plants that, however, likely have a PG layer, this leaves many issues with respect to the composition, exact function, and biosynthesis in land plants to be explored.

## Introduction

Depending on the dating method, plastids emerged between 1.2 to 2.5 billion years ago (Sánchez-Baracaldo et al., 2017; Gawryluk et al., 2019; Strassert et al., 2021; Bowles et al., 2023). An endosymbiotic cyanobacterium was streamlined by a heterotrophic protist of still unknown nature and integrated into its biology, hereby transitioning into the plastid of the archaeplastidal ancestor. From that ancestor, three main phototrophic eukaryotic lineages evolved, the Glaucophytes, Rhodophytes, and Chloroplastida (Strassert et al., 2021; Sibbald and Archibald, 2020). While phylogenomic reconstructions of ancient traits relating to primary endosymbiosis are quite difficult due to large divergence times over evolutionary time, the most recent phylogenomic and network analyses suggest that the plastid donor was a basal-branching freshwater cyanobacterium, whose closest extant sister-lineage is the *Gloeomargarita* genus (Ponce-Toledo et al., 2017). Part of transforming an endosymbiont into an organelle involves transferring the majority of genetic information to the nucleus of the host cell through endosymbiotic gene transfer (EGT) (Deusch et al., 2008). Hence, the vast majority of photosynthesis- and plastid biogenesis-associated genes are cytosolically translated and then imported (Miyagishima et al., 2014; Knopp et al., 2020; Dowson et al., 2022). Enzymes of peptidoglycan (PG) layer biosynthesis, if present, are no exception.

The peptidoglycan polymer provides bacterial cell walls with a structure and rigidity to protect themselves against biotic and abiotic stressors such as osmotic pressure, bacteriophages, heat, and salinity (Vollmer et al., 2008). The PG (or murein) layer is made up of sugars and amino acids, whose biosynthesis is initiated in the cytoplasm of Gram-negative bacteria by the PG (Mur) proteins MurA and MurB that catalyse the formation of UDP-N-acetylmuramic acid (UDP-Mur*N*Ac), which is subsequently appended by the Mur ligase toolkit (consisting of MurC-MurF) to form an UDP-Mur*N*Ac-pentapeptide. Subsequently, the MraY protein attaches this latter pentapeptide to a C55-P complex to produce a C55-PP-MurNAc-pentapeptide, with MurG ligating this pentapeptide to a Glc*N*Ac resulting in the production of C55-PP-MurNAc-(pentapeptide)-GlcNAc. The DDL ligase (D-alanine:D-alanine ligase) catalyses the formation of the D-alanyl-D-alanine dipeptide by fusing two molecules of D-alanine through ATP hydrolyzation (Vollmer et al., 2008; Dowson et al., 2022). The product is fused to the lipid anchor undecaprenol pyrophosphate C55-P by MraY and subsequently flipped into the periplasm via the action of penicillin-binding proteins (PBPs). In the Chloroplastida, PBPs have hitherto only been identified in chlorophytes, streptophyte algae, bryophytes and seedless vascular plants (tracheophytes) (van Baren et al., 2016).

The presence of the PG layer in the plastids of the Archaeplastida has been documented for the Glaucophyta and Chloroplastida, but so far, not for rhodophytes (Björn, 2020). A cyanelle, the glaucophyte plastid, possess a reduced yet still relatively thick PG layer between their two membranes with consequences regarding protein import (Steiner et al., 2005). The PG layer is also present in some members of the green lineage, but the degree to which this trait is conserved in multiple clades remains unresolved (Keeling, 2004; Hirano et al., 2016; Grosche and Rensing, 2017; Li et al., 2020b). Earlier studies established that while peptidoglycan cannot be visualised sometimes via electron microscopy, treatment of plant cells with antibiotics that target the peptidoglycan biosynthetic pathway cause distorted plastid phenotypes (Tounou et al., 2002). In addition, click chemistry in mosses was used to attach an azide-modified fluorophore to the alkyne groups of ethynyl-DA-DA to light up the PG layer in the intermembrane space, which surprisingly confirmed its presence and concentration at the division sites of chloroplasts (Hirano et al., 2016). But while the peptidoglycan layer of mosses has been functionally characterised in parts, the same cannot be said for other embryophytes that possess orthologues for a full PG biosynthetic toolkit.

The increase in the availability of chloroplastidal genome and transcriptome assemblies, coinciding with advances in phylogenomic and bioinformatics methods, have provided new opportunities to study the evolution and diversity of the PG layer in the Chloroplastida. Understanding PG layer evolution in the green lineage that is of great agricultural, ecological, and economic importance (Delwiche and Timme, 2011; Kovak et al., 2022), is important, because: (i) it will help us to better understand how various components of plastid development evolved – for example, when certain differential gene losses and gains occurred – could help explain why some members of the chloroplastida display unique plastid phenotypes (Hirano et al., 2016; Li et al., 2017; de Vries and Gould, 2018; MacLeod et al., 2022), (ii) it could shed light on whether this cyanobacterial relic is a core component of the plastid division machinery (PDVM) of land plants, including some angiosperms, as some data suggest (Homi et al., 2009; Tran et al. 2023). In this study, we undertake a comprehensive and evidence-based phylogenomic approach on 48 genomes from the chloroplastidal supergroup to delineate the distribution of the PG layer in the green lineage. We highlight that genes encoding proteins associated with PG layer biosynthesis have an uncommon distribution in the Chloroplastida and that the PG layer did not evolve concurrently with a component of the plastid division machinery, FtsZ3, as recently suggested (Grosche and Rensing, 2017). The results underscore the PG layer’s existence in gymnosperms and spermatophytes, for which dedicated studies exploring its biological relevance for the plastid organelle are surprisingly sparse.

## Material and methods

### Determining the phylogenetic distribution of PG layer biosynthetic enzymes in the Chloroplastida, and plastid division components in chlorophyte algae

Protein sequence IDs of the ten key enzymes involved in peptidoglycan layer biosynthesis were retrieved from the genome of the liverwort *Marchantia polymorpha* and used as queries for the identification of orthology clusters (Bowman et al., 2017). OrthoFinder version 2.5.4 was used to identify orthologs among the input genomes from 48 Chloroplastida members, with a BLASTp e-value threshold of 1x10^−9^ (supplementary table S1). The phylogenetic distribution of enzymes involved in peptidoglycan layer biosynthesis was determined by examining the presence or absence of orthologous groups (orthogroups) containing these proteins across different members of the Chloroplastida (supplementary table S2). This exact pipeline was replicated to determine the presence/absence of plastid division components in 37 chlorophyte algae and one Prasinodermophyta (Li et al., 2020b; Grigoriev et al., 2021) (supplementary table S3). Furthermore, where a given orthology cluster contained a protein family, which was the case for FtsZ proteins, phylogenetic trees were constructed to separate each protein into a respective subfamily. Sequence alignments were undertaken using MAFFT v7.471 using the LINSI parameter, with tree building being undertaken using IQ-TREE v2.0.3 using an automated selection model, with a 100 non-parametric bootstrap (Katoh et al., 2002; Minh et al., 2020). Finally, we used the SHOOT.bio phylogenetic application to determine whether PG layer biosynthetic genes in some seed plants branch within the terrestrial clade (Emms and Kelly, 2022).

### Phylogenetic species tree construction

The species tree is based on a weighted concatenated alignment from 11 individual alignments. The first step was to calculate protein families including all sequences from the 48 analysed genomes. Pairwise local identities were determined via DIAMOND (v2.0.1) and filtered for all reciprocal best blast hit pairs with at least 40% local sequence identity and a maximum e-value of 1x10^−10^ (Buchfink et al., 2015). A total of 210 protein clusters contained sequences from all 48 genomes, however, no single-copy gene cluster was found. To create a robust reference tree, 11 protein families were chosen in which only few genomes were represented by more than one sequence. For these clusters, alignments were calculated with MAFFT v7.471 using the LINSI parameter and the duplicate sequences were manually removed, favouring the copies that did not show major deletions or insertions to yield a robust phylogeny (Katoh et al., 2002). All 11 alignments were concatenated while equalizing their phylogenetic signal using a weighted concatenation approach. The final tree was built by IQ-TREE v2.0.3 (Minh et al., 2020) with 100 non-parametric bootstraps using the LG+F+R7 substitution model. Best-fit model identification via IQ-TREE’s model finder (Minh et al., 2020). Tree trimming and visualization was carried out using the ggtree R package (Yu et al., 2017). A species tree of chlorophytes – used to plot the phylogenetic distribution of plastid division machinery components in this phylum – was estimated using STAG in the OrthoFinder run (Emms and Kelly, 2015, 2015, 2018).

### Delineating domain architecture and function of orthologous sequences

InterProScan v5 (Jones et al., 2014) was used to delineate the basic domain architecture and function of protein sequences involved in peptidoglycan layer biosynthesis. The program was used to identify protein domains, annotate their functions, and determine the arrangement and composition of the domains in the protein sequences.

## Results

### Phylogenetic distribution of PG layer biosynthesis genes across the Chloroplastida

The green lineage is comprised of three clades: Streptophyta, Chlorophyta and the basal-branching Prasinodermophyta (Li et al., 2020b) (Fig. 1). The Chlorophyta and Prasinodermophyta are comprised entirely of algae, whereas the Streptophyta are comprised of algae and embryophytes (land plants). While the distribution of a full PG layer biosynthetic toolkit is mostly concentrated and noticeable in streptophyte algae, bryophytes (non-vascular plants) and lycophytes, this toolkit is also present in the Prasinodermophyta species *Prasinoderma coloniale* and four phylogenetically distant chlorophyte species: *Micromonas pusilla* (Mamiellophyceae), *Chloropicon primus* (Chloropicophyceae), *Ulva mutabilis* (Ulovophyceae) (Fig. 1). The protein domain and gene ontology analyses show a high level of conservation and confirms that these proteins likely play key roles in peptidoglycan biosynthesis (supplementary table S4).

**Fig. 1:**
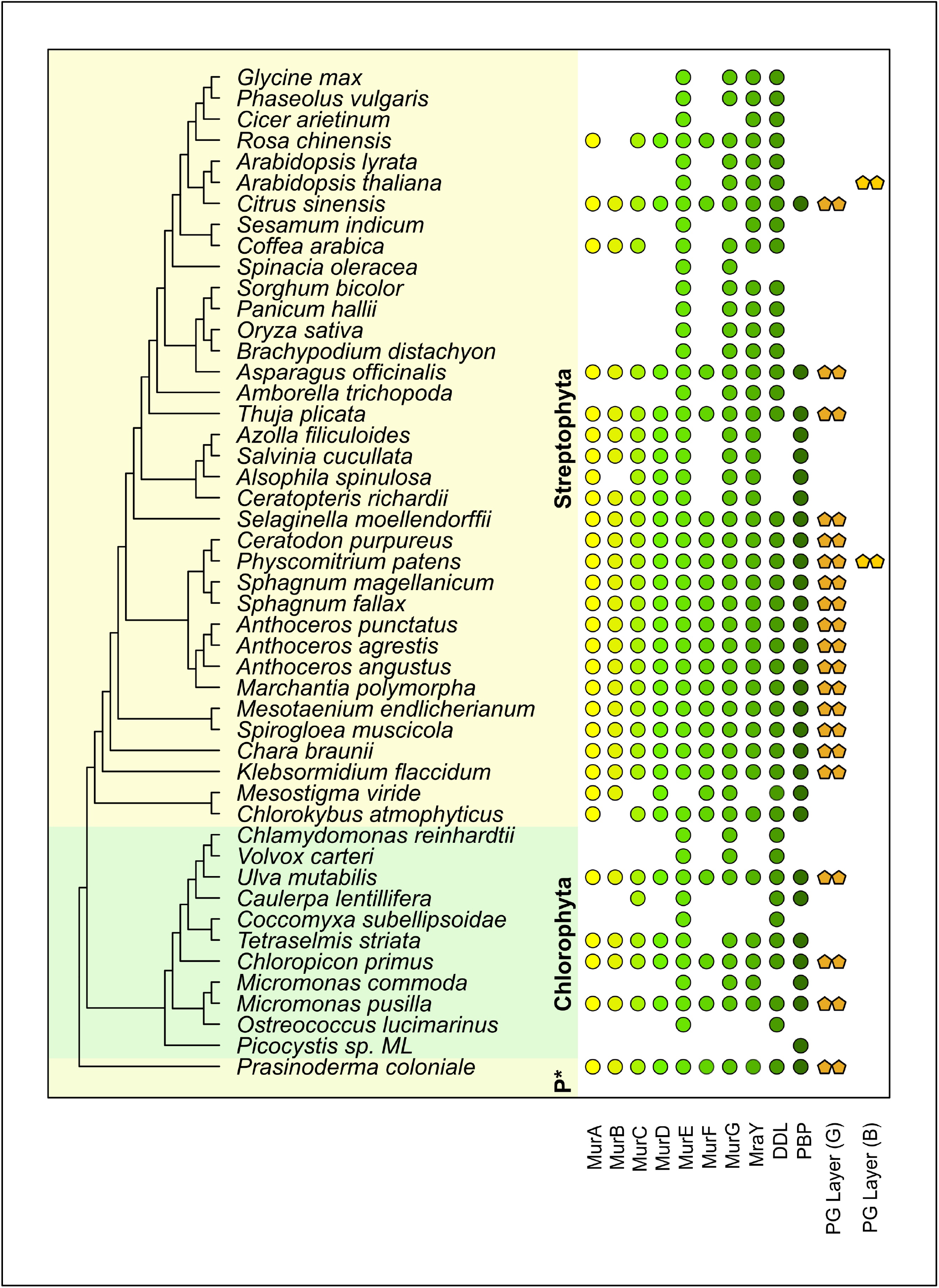
Phylogenetic distribution of the PG layer biosynthetic toolkit across the Chloroplastida. The likely presence of a PG layer is indicated by the identification of all enzymes involved in its biosynthesis. Note that there are cases where not all enzymes of the pathway can be identified through bioinformatic means (indicated by *PG Layer (G)*), although experimental click chemistry, adding an azide-modified fluorophore to the ethynyl-DA-DA of the PG layer, indicates its presence (*PG Layer (B)*). *P\**; Prasinodermophyta.

Furthermore, while previous studies have suggested that the PG layer was differentially lost in the MRCA of seed plants (spermatophytes) (van Baren et al., 2016; Grosche and Rensing, 2017), the full toolkit for the biosynthesis of peptidoglycan is identified in at least three phylogenetically distant members of the seed clade: *Thuja plicata* (Gymnosperms), *Asparagus officinalis* (Monocots), and *Citrus sinensis* (Eudicots) (Fig. 1). This includes the identification of PBP family members in spermatophytes (Fig. 1 and 2). In addition, there is strong structural conservation in the PG layer biosynthetic toolkit with respect to proteins and domain architecture, and from algae to angiosperms (Jones et al., 2014) (Fig. 2).

**Fig. 2:**
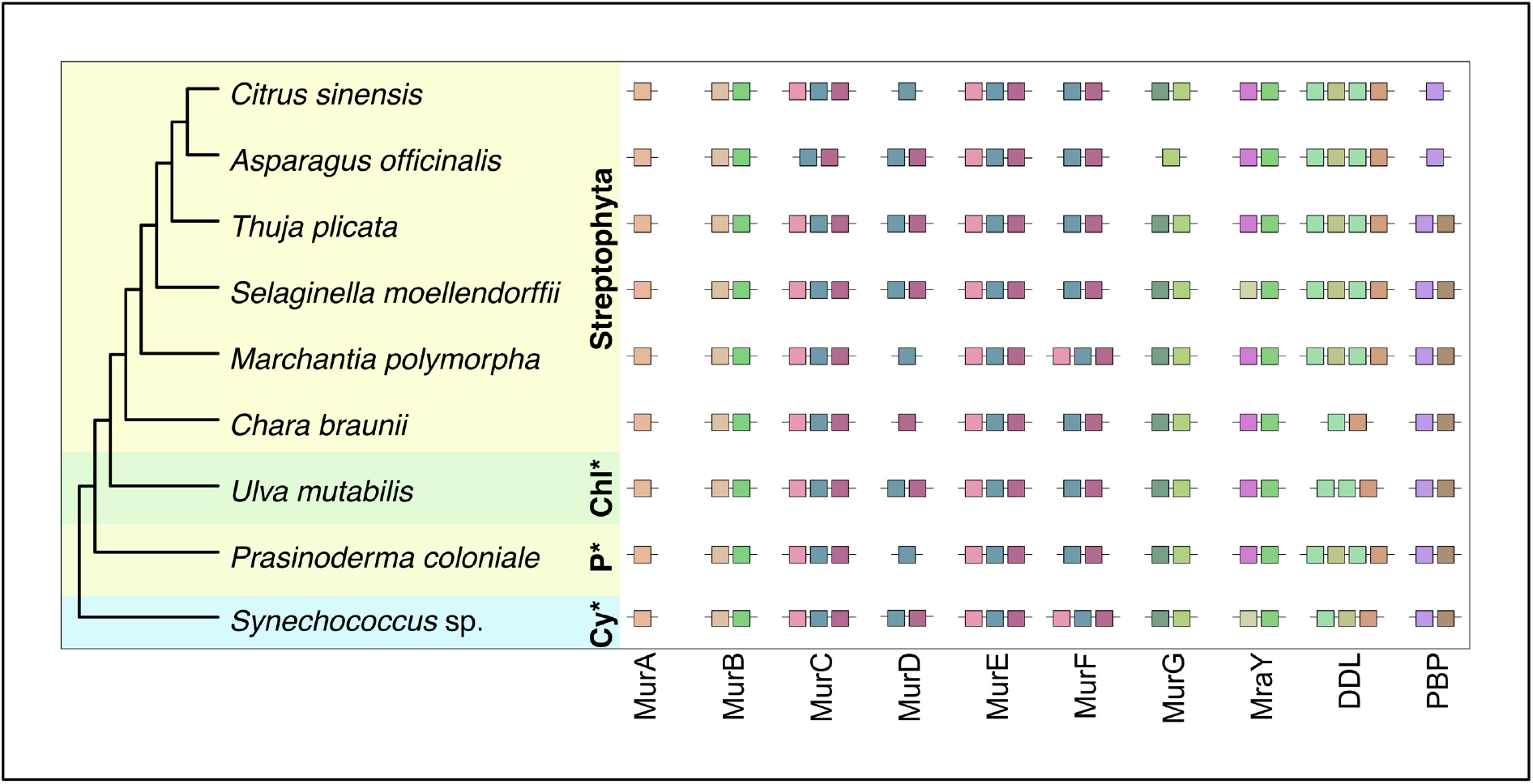
The PG layer biosynthetic toolkit is structurally well conserved from cyanobacteria to angiosperms. *Cy\**, Cyanobacteria; *P\**, Prasinodermophyta; *Chl\**, Chlorophyta. A full key corresponding to each domain’s identity can be accessed in supplementary Fig. S1.

### No evident correlation between the presence of the PG layer and any one of the three FtsZ proteins

The PG layer plays a key role in regulating chloroplast division in bryophytes and streptophyte algae (Machida et al., 2006; Homi et al., 2009; Hirano et al., 2016; Grosche and Rensing, 2017; Dowson et al., 2022). The GTPases FtsZ1 and FtsZ2, forming a dynamic heteropolymer, are involved in regulating the formation of the inner plastid division ring, or Z-ring (Yoshida et al., 2016). FtsZ3 also likely plays a crucial role in Z-ring formation, via similar methods as its two FtsZ cousins (Martin et al., 2009), but is absent from many genomes (Fig. 3). The dynamin-like ARC5 protein polymerizes on the chloroplast’s outer surface, leading to the formation of an outer division ring (Gao et al., 2003), where FtsZ3 is likely involved, although it is unclear whether it associates directly with ARC5 (Martin et al., 2009).

**Fig. 3:**
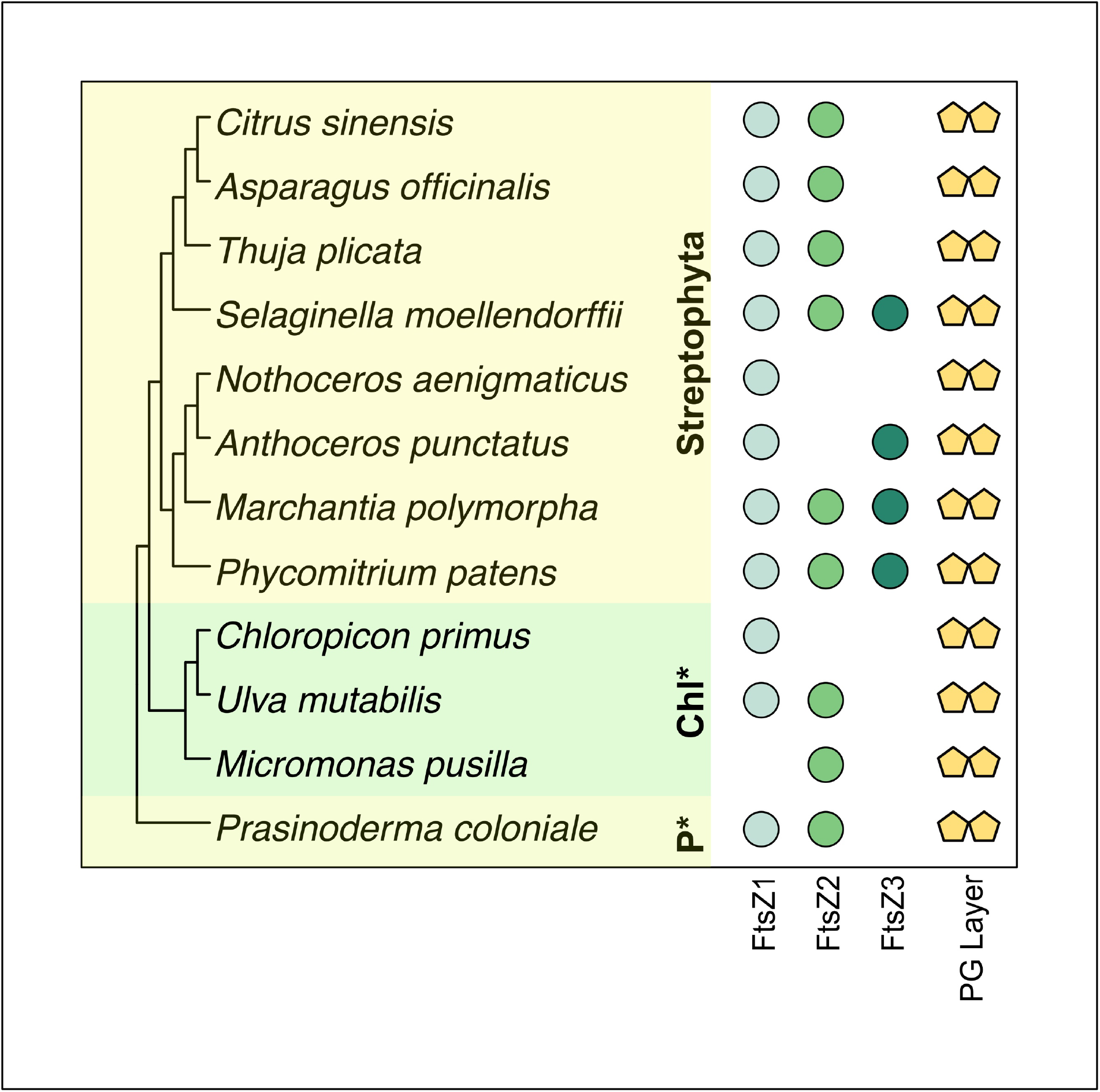
Phylogenetic distribution of the FtsZ plastid division proteins in Chloroplastida that likely have a plastid enveloping PG layer, highlighting the unlikely coevolution between FtsZ3 and the PG layer. *P\**, Prasinodermophyta; *Chl\**, Chlorophyta. Orthologue metadata for FtsZs in the Streptophyta was obtained from MacLeod *et al*. 2022.

FtsZ3 was suggested to play a role in regulating the biogenesis of the PG layer due to an alleged correlation between these two traits (Grosche and Rensing, 2017). There are, however, multiple exceptions to this correlation. For example, *Thuja plicata, Asparagus officinalis* and *Citrus sinensis* possess orthologues representing a full enzymatic toolkit for PG layer biosynthesis (Fig. 1 and 2), but lack FtsZ3 (Fig. 3). In addition, the hornwort *Nothoceros aenigmaticus* and the chlorophytes, *C. primus, U. mutabilis*, and *M. pussilla* all likely possess a PG layer between their chloroplast membranes (supplementary Fig. S2), but lack FtsZ3. In summary, the now-available genomes do not support a PG layer and FtsZ3 co-evolution or functional connection. In fact, it appears that the presence of the chloroplast PG layer is not dependent on the presence of any one specific protein of the FtsZ family (Fig. 3).

## Discussion

Data emerging from both the screening of new genomes becoming available and biochemical characterizations shows that a PG layer is found enveloping the chloroplasts of more taxa, including some angiosperms, than previously thought. The presence of a PG layer has been explicitly mentioned in previous studies for only a few green algae such as *M. pusilla, Picocystis salinarum* (Picocystophyceae), and *Nephroselmis pyriformis* (Nephrophyceae) (van Baren et al., 2016). We highlight the presence of a full suite of PG layer synthesizing enzymes in several additional algae (*C. primus, U. mutabilis* and *T. striata*) and underscore the data available for many land plants, now including proteins of the PBP family, that likely possess this ancient trait. Unusual phylogenetic distributions of genes and entire pathways are sometimes the result of the identification of bacterial false positive contaminations (Koutsovoulos et al., 2016; Husnik and McCutcheon, 2018; Goig et al., 2020), which is unlikely with respect to the enzymes of PG layer biosynthesis considering the robust branching of, for instance, *T. plicata, A. officinalis* and *C. sinensis* deep within the embryophyte clade (supplementary table S5). That is, all available data suggest a monophyletic origin of the pathway (Li et al., 2020b) (Fig. 1) and an independent differential loss in various taxa across the Archaeplastida.

Most angiosperms do not seem to encode for a complete set of enzymes synthesizing the PG layer. They all share, however, four enzymes related to the process (MurE, MurG, MraY, and DDL), called the “4-PGN” set (Fig. 1) and recent experimental work suggests that two angiosperms, *Arabidopsis thaliana* and *Nicotiana benthamiana*, may have a PG layer surrounding their chloroplasts (Tran et al., 2023). If true, then it would suggest that these species use a different set of enzymes and biochemistry to synthesize parts of the PG layer, with the 4-PGN set playing a key role, therefore being retained. Intriguingly, the retention of the same set of genes (+/-1) occurred independently in some chlorophyte algae such as *Micromonas commoda* (Fig. 1), raising the question whether they have been retained for the same functional reason.

While recent biochemical and metabolomic analyses suggest that components of the moss peptidoglycan biosynthetic pathway – specifically, the active sites of core ligase enzymes – display strict conservation in comparison to the PG layer biosynthetic pathway of cyanobacteria (Dowson et al., 2022), the FtsZ3 PDVM component is unlikely to play a role in PG layer biosynthesis in moss. Genome analyses indicate that the PG layer exists in all three phyla of chloroplastida (van Baren et al., 2016; Grosche and Rensing, 2017; Li et al., 2020b), including our own analysis. There is, however, no strict connection between FtsZ3 or any FtsZ gene and this cyanobacterial relic in terms of the FtsZ-based ring’s association with the PG layer. Therefore, any gene from the FtsZ family can likely perform its role in regulating the formation of Z- and outer rings, indicating functional redundancy within this family.

## Conclusion

The peptidoglycan layer of chloroplasts was present in the MRCA of Chloroplastida and lost in most Chlorophyta and many Streptophyta, but retained in the Prasinodermophyta. Since the number of annotated genome assemblies for this latter phylum still stands at a mere one, however, it will be interesting for future genome mining experiments to elucidate whether the PG layer can be characterized – either biochemically or genomically – in this basal-branching green phylum. One can conclude that the PG layer is present in the chloroplasts of at least three spermatophytes, likely more, and from what we can tell our screening is the first to identify the presence of the PBP family in this clade. In addition, we bring into the spotlight two species of chlorophytes whose chloroplasts are likely enveloped by a PG layer, in summary suggesting that peptidoglycan is more widespread in the chloroplasts of this phylum than previously thought, while there is no longer a support regarding the correlation between the presence of the PG layer and the plastid division protein FtsZ3. With this evidence at hand, studies now need to elucidate both the biochemical nature as well as the biological relevance of the PG layer in angiosperms that were, until recently, thought to lack this ancient cyanobacterial trait.

## Data availability

All data generated in this study can be accessed here: https://uni-duesseldorf.sciebo.de/s/oydhtETq041Keop

## Acknowledgements

We thank Professor William F. Martin for his ongoing support.

