## Supplementary Figures S1-S2 for "A mysterious cloak: the peptidoglycan layer of algal and plant plastids"

All data generated in this study can be accessed here:
https://uni-duesseldorf.sciebo.de/s/oydhtETq041Keop

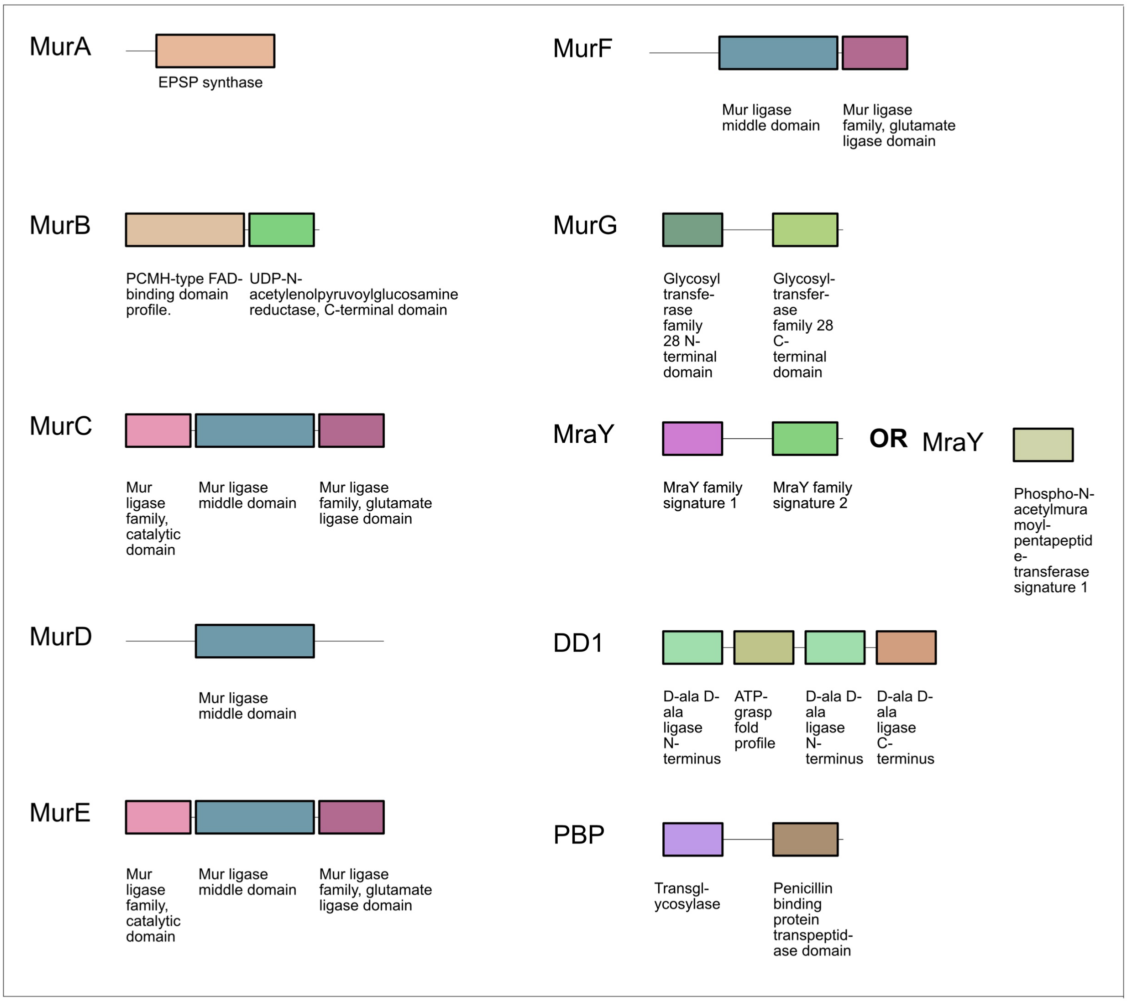


***Supplementary figure S1.*** Domain annotations of the murein layer biosynthetic toolkits from *Prasinoderma coloniale* and *Selaginella moellendorffii*

All data generated in this study can be accessed here:
https://uni-duesseldorf.sciebo.de/s/oydhtETq041Keop

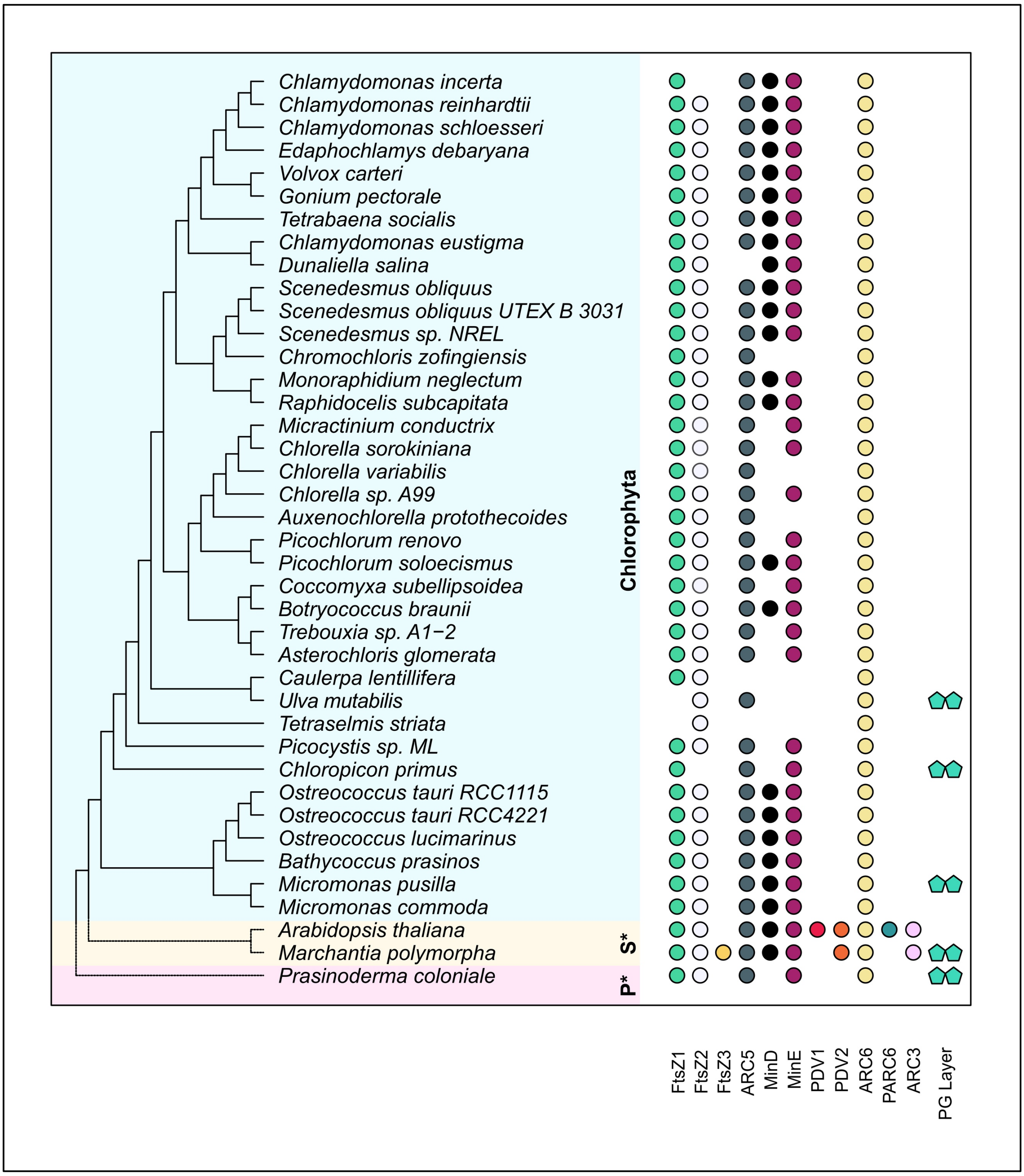


***Supplementary figure S2.*** **Phylogenetic distribution of plastid division machinery components in chlorophytes and prasinodermophytes.** *S**, Streptophyta; *P**, Prasinodermophyta.
